# High Bandwidth Power Electronics and Magnetic Nanoparticles for Multichannel Magnetogenetic Neurostimulation

**DOI:** 10.1101/2021.06.23.447876

**Authors:** Boshuo Wang, Zhongxi Li, Charles E. Sebesta, Daniel Torres Hinojosa, Qingbo Zhang, Jacob T. Robinson, Gang Bao, Angel V. Peterchev, Stefan M. Goetz

**Affiliations:** Department of Psychiatry and Behavior Sciences, School of Medicine, Duke University, Durham, NC 27710, USA; Department of Electrical and Computer Engineering, School of Engineering, Duke University, Durham, NC 27708, USA; Department of Bioengineering, Rice University, Houston, TX 77005, USA; Department of Electrical and Computer Engineering, Rice University, Houston, TX 77005, USA; Department of Neuroscience, Baylor College of Medicine, Houston, TX 77030, USA; Department of Neurosurgery, School of Medicine, Duke University, Durham, NC 27710, USA; Department of Biomedical Engineering, School of Engineering, Duke University, Durham, NC 27708, USA

**Author notes:** These authors contributed equally to the work (Boshuo Wang, Zhongxi Li).

**Keywords:** magnetogenetic neurostimulation, frequency-division-multiplexing, alternating magnetic field, GaN transistors, multichannel stimulation, power electronics

## Abstract

**Objective:** We present a power electronic system and magnetic nanoparticles for multiplexed magnetogenetic neurostimulation with three channels spanning a wide frequency range and rapid channel switching capability. This enables selective heating of magnetic nanoparticles with different coercivity using various frequency–amplitude combinations of the magnetic field. Such multiplexed operation could provide the technical means for selective magnetogenetic neurostimulation beyond its spatial focality limits.

**Approach:** The electronic system uses a hybrid of silicon metal–oxide–semiconductor and gallium-nitride field-effect transistors, which generate the required high-amplitude current up to 500 A in the sub-MHz range and the high-frequency current in the MHz range, respectively. Via three discrete resonance capacitor banks, the system generates an alternating magnetic field in the same liquid-cooled field coil at three distinct frequency channels spanning 50 kHz to 4 MHz. Fast switching between channels is achieved with high-voltage contactors connecting the coil to different capacitor banks. We characterized the system by the output channels’ frequencies, field strength, and switching time, as well as the system’s overall operation stability. Three types of iron oxide nanoparticles with different coercivity are developed to form three magnetothermal channels. Specific absorption rate and infrared thermal imaging measurements are performed with the nanoparticles to characterize their heating and demonstrate selective actuation for all three channels.

**Main results:** The system achieved the desired target field strengths for three frequency channels (70 kA/m at 50 kHz, 10 kA/m at 500 kHz, and 1 kA/m at ≥2 MHz), with rapid switching speed between channels on the order of milliseconds. The system can operate continuously for at least two hours at 30% duty cycle with 125 Arms load in the coil, corresponding to a stimulation protocol of cycling the three channels at target strength with 3 s pulses and 7 s interpulse intervals. The nanoparticles were heated with selectivity between 2.3× and 9× for their respective frequency channel. The system’s intended use was thus validated with three distinct channels available for magnetothermal heating.

**Significance:** We describe the first combination of a power electronic system and magnetic nanoparticles that achieves three stimulation channels. Selective actuation of nanoparticles is demonstrated for each channel using the same field coil, including a novel composition responding to magnetic fields in the MHz range. This approach could improve the speed and flexibility of frequency-multiplexed magnetogenetic neural stimulation.

## 1. Introduction

Alternating magnetic fields serve for a wide range of applications in medicine and biological research. Strong alternating magnetic fields applied to magnetic nanoparticles can generate heat at the location of the nanoparticles at depth within biological tissue. This local heating can be used for hyperthermal cancer treatment (Hayashi *et al.* 2010; Shaterabadi, Nabiyouni and Soleymani 2018), genetic transfection (Dahmani *et al.* 2013), protein manipulation (Monsalve *et al.* 2015), or site-specific drug release (Oliveira *et al.* 2013; Wang *et al.* 2016). When the magnetic nanoparticles are in close proximity or bound to cells expressing temperature-sensitive transient receptor potential (TRP) channels, it is possible to stimulate neural activity in a genetically targeted population. This process is known as magneto-thermal genetics or magnetogenetics (Huang *et al.* 2010; Chen *et al.* 2015; Munshi *et al.* 2017; Duret *et al.* 2019). Although mechanical forces of magnetic fields on the nanoparticles connected to the channels might play a role, local heating on the molecular level is currently considered more relevant (Christiansen, Senko and Anikeeva 2019). While there have been reports of magnetogenetics based on biogenetic ferritin nanoparticles, the mechanism of action remains unclear as the heat generated by these weakly magnetic nanoparticles is likely to be insufficient to activate thermoreceptors (Stanley *et al.* 2012; Meister 2016; Wheeler *et al.* 2016).

With the ability to activate genetically targeted cell types, magnetogenetic neural stimulation could be a breakthrough for minimally invasive neuromodulation. Transcranial electrical (TES) and magnetic stimulation (TMS) have natural focality limits for the field they form about 15 millimeters below the skull (Dmochowski *et al.* 2011; Goetz and Kammer 2021). In addition to the distance effect, the transcranially injected current of TES spreads out as nontarget areas conduct similarly well and due to current-shunting effects of skull openings (e.g., the eyes) (Bortoletto *et al.* 2016; Guler *et al.* 2016). The near field of magnetic coils without any further selectivity mechanisms as used in TMS similarly spreads out and can in principle not be focused in depth if the coil–target distance is below the wavelength (Deng, Lisanby and Peterchev 2013; Gomez, Goetz and Peterchev 2018). Magnetogenetic stimulation, in contrast, can tune the receiver to achieve practically unlimited focality and selectivity also in the magnetic near-field. Compared to conventional TES or TMS, magnetogenetics benefits from both the *transparency* of tissue to magnetic fields as well as the focality and/or selectivity of the stimulation effect due to the localization of the magnetic nanoparticles and/or selective expression of ferritin (Stanley *et al.* 2012).

In addition to a localization of sensitive ion channels, frequency separation of magnetic effects has been achieved via variation of the size and doping of magnetic nanoparticles (Moon *et al.* 2020; Sebesta *et al.* 2021). Thus, multichannel stimulation could allow high selectivity even between closely located neural elements. In addition, the magnetic field can still be focused with conventional methods to separate targets with wider spacing as in TMS. However, for multichannel stimulation with frequency-selective ion channels, the field source needs to be able to generate widely different frequencies at various amplitudes. Reasonable signaling and driving of network effects requires short turn-on and turn-off times of the field and rapid changes between frequency channels.

Generating alternating magnetic fields has been a challenging laboratory effort mainly due to the high field strength required. These high field strengths are a consequence of typically extremely weak electromagnetic coupling between the magnetic source (e.g., coils) and the target so that most energy stays with the source. Thus, powerful power electronic sources are typically required to deliver at least couple of hundred amps at relative high frequency (e.g., at hundreds of kilohertz) to a coil. The required current amplitude and frequency often overwhelm the common power electronics portfolio and call for significant trade-offs between power, cost, and the strength of the field versus the frequency and scale of the field (Christiansen *et al.* 2017). Most methods for generating magnetic fields are limited to the kilohertz range due to the use of ferrite cores in the field coil. Although in-vitro characterization of nanoparticles has been performed with low power in the megahertz range (e.g., several MHz to 40 MHz) (Huang *et al.* 2010; Davis *et al.* 2020), the miniaturized field coils limit the spatial extent of the field significantly for application in vivo, where the targets are located *outside* the coil. Achieving the required field strength at distance and scale for megahertz frequency is rather challenging for the power electronics because of high-frequency effects such as proximity as well as skin effects, parasitic inductances, capacitances, and coupling.

Prior research often uses general linear amplifiers with power levels of several hundred watts to drive the magnetic coil (Huang *et al.* 2010; Rohan *et al.* 2014). The advantage of general-purpose amplifiers is the relatively wide band and continuously adjustable frequency. However, general-purpose linear amplifier circuits have limitations in efficiency, power, and frequency. Known amplifier circuits, most commonly of classes A, B, AB, or C, typically do not exceed 50% and can for reasons of principle not exceed ∼75% efficiency in their linear range and most of the time operate notable further below that limit (Hilbourne and Jones 1955). Paralleling more transistors does not solve the problem as they operate as controllable resistance and necessarily generate loss to carve the AC output from their DC supply. For strong fields, the low efficiency in combination with the necessary high current rapidly heats up the amplifier and limits the power. For typical currents up to 500 A, the consequence would be excessive requirements for cooling and the necessary power supply. Furthermore, the bandwidth of conventional high-power amplifiers is limited for several reasons. First, parasitic capacitances and inductances limit the speed of high-power circuits. Second, the electrical impedance associated with the coil inductance depends on the frequency. When the field frequency is changed by a factor of 100 (50 kHz to 4 MHz here) for good separation between several channels, the necessary voltage likewise increases by a factor of 100 for approximately equal power. Conventional amplifier circuits do not provide a solution for this problem.

A common way to produce RF fields leverages L–C resonance, where energy oscillates between a coil (L) and a capacitor (C). In this case, the power electronics does not have to deliver reactive power to the coil, but only feeds the losses due to internal resistance, field absorption, and wave emission. The resonant circuit furthermore compensates the majority of the inductive impedance of the coil so that for series resonance the necessary voltage for the power electronics does no longer grow linearly with the frequency into unacceptable ranges (for parallel resonance, the current could be limited). Thus, resonant circuits provide a relatively *effortless* way for strong magnetic fields since already a low driving voltage is capable of producing high currents. Resonant circuits are known from more remote applications, such as transcranial magnetic stimulation or inductive charging (Peterchev, Deng and Goetz 2015; Guidi *et al.* 2017).

However, series-resonant circuit technology according to the state of the art entails several limitations. First, the output frequency of a resonator is widely fixed. We will solve this problem by implementing multiple sets of capacitors with rapid software-controlled multiplexing in between. High-current, high-voltage contactors are used for multiplexing; the switching time is on the order of milliseconds to satisfy most applications whereas the added conduction loss is minimal. Second, despite not having rapidly growing voltage, the wide frequency range of more than six octaves required here is challenging due to the high currents pushing conventional silicon (Si) transistors to their limits. At low frequencies, the power electronics have to drive up to 500 A, requiring high-power silicon transistors. While these high-power silicon transistors serve well for the low frequency range (kHz), they exhibit large parasitics, such as the gate and the drain–source capacitance, and thus struggle in the MHz range (Hilbourne and Jones 1955).

Considering these problems, we propose a hybrid silicon–gallium-nitride field-effect transistor (FET) circuit with switching modulation and adaptive series resonance. Compared to linear amplifiers, switched circuits deliver poor output quality with harmonic content, but the resonant configuration serves as a sharp filter and only lets sinusoidal current pass. The hybrid configuration with Si and gallium-nitride (GaN) exploits the advantages of each technology while avoiding their issues: vertical trench Si FETs with their high-power but limited speed due to parasitics and low carrier dynamics deliver the majority of power from low to medium frequencies (50 kHz – 500 kHz); lateral GaN-on-Si FETs with their high speed but limited thermal and current capabilities take over at high frequencies (>500 kHz). We demonstrate a system with three channels (Table 1), each separated by a factor of ten in frequency and field strength for the envisioned amplitude-frequency product (Hergt and Dutz 2007; Christiansen *et al.* 2017)

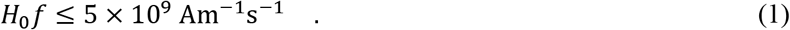

**Table 1.** Design specification of the three channels with nominal value and target range in parentheses.

| Parameter | Units | Channel 1 |  | Channel 2 |  | Channel 3 |  |
| --- | --- | --- | --- | --- | --- | --- | --- |
| Frequency | kHz | 50 | (50–100) | 500 | (380–580) | 5000 | (2000–5000) |
| Magnetic field | kA/m | 70 | (50–100) | 10 | (8–12) | 1 | (0.8–1.6) |
|  | mT | 88 | (63–125) | 13 | (10–15) | 1.3 | (1–2) |
| Amplitude-frequency | $10^9 \cdot \text{Am}^{-1}\text{s}^{-1}$ | 3.5 | (3.5–5) | 5 | (3–5) | 5 | (4–5) |

These three channels were chosen to demonstrate the possible wide range of frequencies, covering existing magnetic nanoparticles with good selectivity (Moon *et al.* 2020; Sebesta *et al.* 2021) and exploring novel nanoparticles responding preferably to higher frequencies. The concept scales well and should allow further channels. Smaller frequency differences can be built by combination of the implemented capacitor options. Notice that the product *H*_0_*f* is constant across the three frequencies, thus ensuring a similar stimulus effect in either channel. As a result, we would expect higher current at low frequency and lower current at high frequency, which respectively perfectly fits into either Si or GaN FETs’ operating profiles. The controller can drive subsets of the transistors separately to adaptively mix the transistor technologies and reduce the number of driven transistors when the gate loss starts dominating at higher frequencies.

## 2. Methods

### 2.1. Operating principle and system overview

We employ series L–C oscillation to produce alternating current in the coil (*L*_coil_ = 3.14 μH, see Section 2.5) to generate the alternating magnetic field. In our system, we specifically attach three capacitor banks to the same coil, with each bank properly tuned to achieve the desired oscillation frequency. The design allows simple expansion to additional channels. For each L–C-tank combination, the hybrid full-bridge circuit generates rectangular voltage output with adjustable duty cycle and frequency controlled by the system-on-chip (SoC) controller. The series resonance as a sharp notch-filter presents a large impedance to all frequency contents of the rectangular driving voltage except for its fundamental so that a sinusoidal current follows. Whereas the duty cycle controls the coil current’s amplitude, the frequency must match either resonance frequency of the L–C bank to excite it, which can be easily tuned on the software side. Once the setup is tuned, the driving circuit only needs to operate at a low voltage (here ±36 V peak for the rectangular waveform) and the resonance design shields off the reactive component. Thus, the power electronics mostly feeds the power loss and field emission of the oscillator. Despite the low voltage at the power-electronics side, the coil and capacitor voltages can reach 1 kV rms. On the flip side, the coil current (up to 500 A rms for Channel 1 – Ch 1) must entirely flow through the circuit, requiring proper wiring and cooling design.

The actual implementation extends from the above description in two aspects: 1) two driving circuits of different semiconductor technologies are used, each optimized for one or two operating frequencies; 2) we use contactors to decouple different channels.

### 2.2. Implementation of electronics

Instead of one transistor technology for the entire range of frequencies and currents, two specialized device types work collaboratively for various trade-offs (Figure 1): for Channel 3 (Ch 3), the power switches of the circuit needs to operate on the order of *f* ≥ 2 MHz but only conduct low current (e.g., *I*_coil_ ≤ 10 A rms). Latest high-electron-mobility GaN-type transistors appear to fit this scenario perfectly, due to their lower parasitic capacitance at gate and output, shorter carrier lifetimes, higher critical field strength, and higher carrier mobility (Zhang *et al.* 2012; Meneghesso, Zanoni and Meneghini 2014). On the other hand, the operating current of Ch 1 can reach beyond 500 A. Vertical Si power trench FETs are more suitable here due to their large die areas.

**Figure 1.**
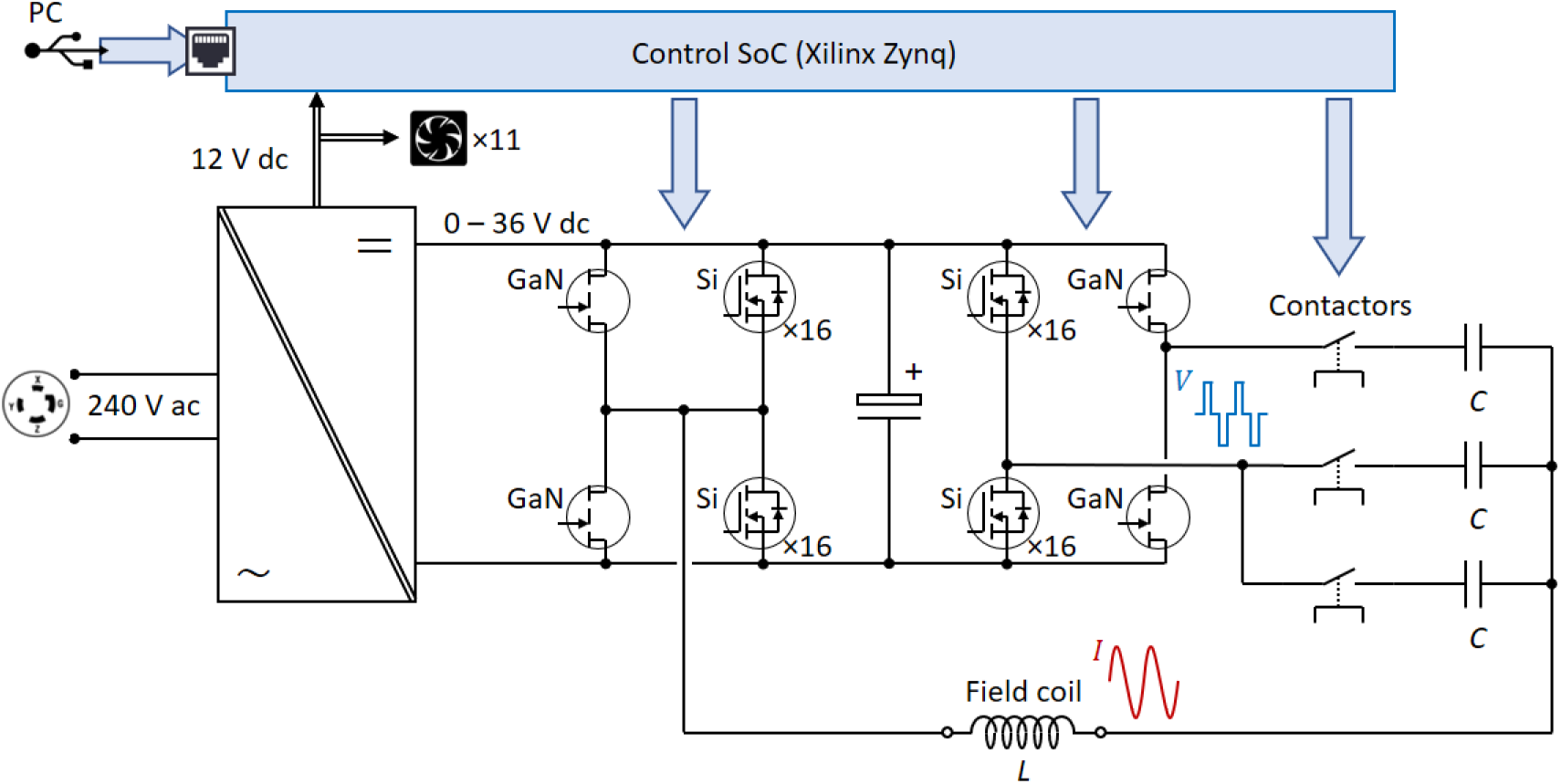
The system overview of the electronics. The control signals are denoted with blue arrows.

The electronic circuit is modularized, with each module sharing a similar layout for better scaling. The full bridge circuit for the 50 kHz channel is implemented with 16 FETs (IPT015N10N5, Infineon Technologies AG, Neubiberg, Germany) for each switch due to the peak current demand. At 500 kHz, the controller activates four FETs per switch at a time in a round-robin fashion to reduce the switching loss. As for the MHz channel, the driving voltage is provided by a full bridge with four GaN transistors (GS61008T, GaN Systems, Ottawa, Canada).

Finally, three high-voltage contactors (B88269X1090C011, TDK Electronics AG, Munich, Germany) determine which channel is activated. The contactors avoid circulating currents between different channels and allow the magnetic coil to be electrically connected to one set of the corresponding series capacitors for each channel at a time, with switching on milliseconds timescale. The contactors are driven by MOSFETs (ZVN4210A, Diodes Incorporated, Plano, TX, USA) via the SoC (see Section 2.5). Table 2 lists the major hardware configuration of the electronics.

**Table 2.** List of major components and technical specification of the three channels.

|  | Channel 1 | Channel 2 | Channel 3 |
| --- | --- | --- | --- |
| Semiconductor technology | Si, vertical trench |  | GaN, planar |
| Transistor | IPT015N10N5 |  | GS61008T |
| Contactors | B88269X1090C011 |  |  |
| DC power supply | 36 V, 1 kW×5 | | |
| Operation frequency | 50 kHz | 500 kHz | $\geq 2$ MHz |
| Coil current (rms) | $\leq 500$ A | $\leq 100$ A | $\leq 10$ A |
| Coil voltage(rms) | $\leq 600$ V | $\leq 700$ V | $\leq 900$ V |
| Peak magnetic field | 100 mT | 20 mT | 4.5 mT |
| Peak field power | 155 kVA | 33 kVA | 3 kVA |
| Peak active (driving) power | 4.5 kW | 100 W | 1 W |

### 2.3. Resonance capacitors and cable

The capacitor bank consists of a circuit board (Figure 2, left) with film capacitors of various types and sizes (Cornell Dubilier Electronics, Liberty, SC, USA and WIMA GmbH & Co. KG, Mannheim, Germany). Quick-change sockets (SparkFun Electronics, Niwot, CO, USA) are used so that capacitors can be easily switched in and out to adjust the capacitance of the resonance circuit and thus the frequency of each channel (Figure 2, left and right). As the power electronics’ output frequency is limited by the FPGA’s clock frequency (100 MHz), the adjustment in the megahertz range is more granular and the output frequency of Ch 3 might not meet the exact resonance frequency of its L–C circuit using discrete capacitors. Therefore, a variable capacitor (10–100 pF, Sprague Goodman Electronics, Westbury, NY, USA) was installed in series with the replaceable film capacitors, which allows the user to tune the circuit to resonate at the discrete frequencies output in the MHz range and maximize the output current and magnetic field. For Ch 1 and Ch 2, the frequency step size is fine enough. Thus, resonance can be reached by adjusting the output frequency via the user interface (see Section 2.5). Care must be taken to check the voltage rating of the film capacitors since the 36 V DC input misrepresents the peak voltage (up to 1 kV) across the capacitors and inductors.

**Figure 2.**
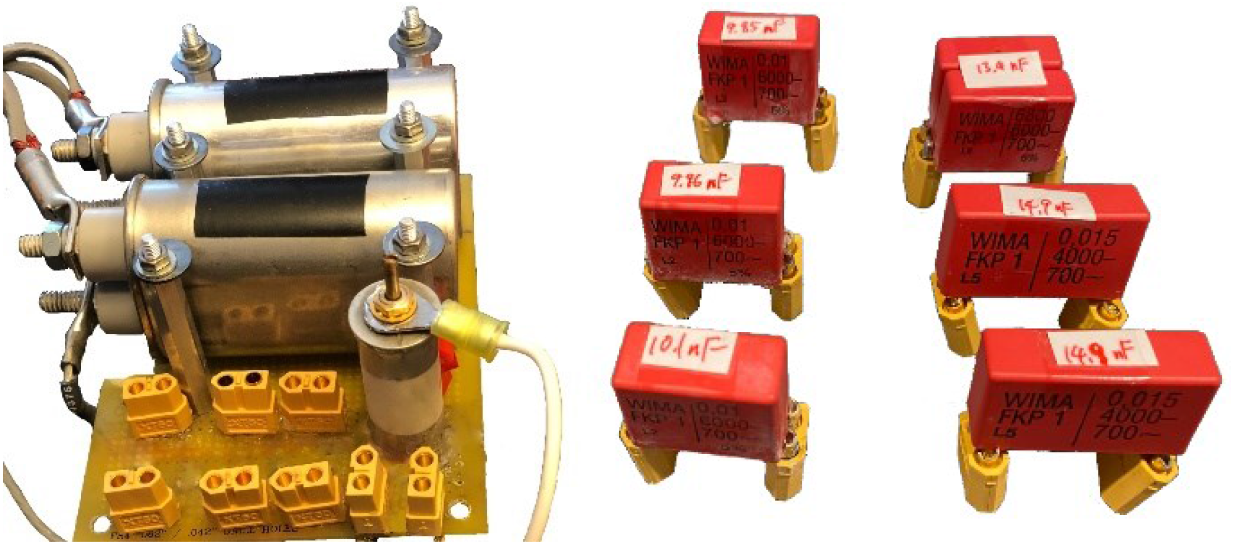
Capacitor bank of the system. Left: capacitor board with sockets (yellow) for interchanging the resonance capacitors for Ch 2 and Ch 3 and the variable capacitor (grey cylinder) for Ch 3. Center: capacitors on quick-change sockets for adjusting the resonance frequency of Ch 2 and Ch 3. Right: Layout of the Litz wires within the cable, with the interference shielding as the outer-most layer.

The electrical cable of approximately 30 cm length (see Figure 5, right) consists of six interleaved 12-AWG type-8 Litz wires (New England Wires Technologies, Lisbon, NH, USA), with three wires in each direction. The cable is ensheathed in interference-shielding heat-shrink tubing to reduce radio-frequency noise leakage and terminated in compression lugs that connect to the coil (Figure 3, left).

**Figure 3.**
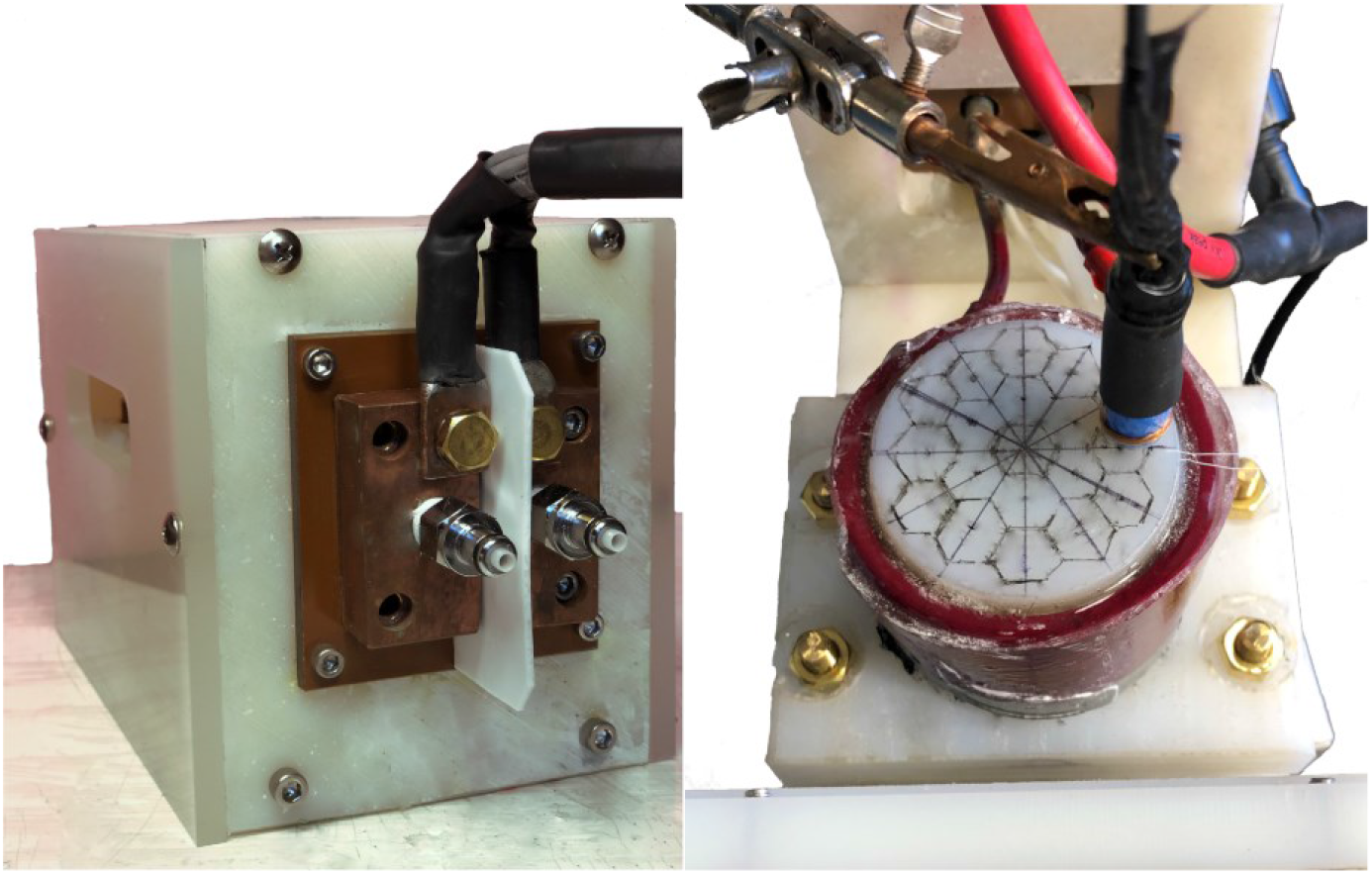
Magnetic field coil. Left: The MSI coil’s custom-requested electric and circulation interface, showing the cable (black) attached to the two ¼-20” bolts and the two quick-disconnect fitting. Right: The top and side panels of enclosure are removed to show the coil windings potted around the central cylinder with a hexagonal grid for mapping the magnetic field; the custom-made field probe (blue and black) is placed against the surface of the cylinder and a Rogowski current waveform transducers (red) measures the coil current.

### 2.4. Magnetic Stimulation Coil

The magnetic field coil (Hi-Flux module, MSI Automation, Wichita, KS, USA) consists of a 6-turn solenoidal coil of 5.75 cm inner diameter and 3.8 cm height wound with a 3/16-inch copper tube, which allows liquid cooling with water (Figure 3). The coil contains a ferrite core (μ = 2300, Ferroxcube 3C90, Eindhoven, Netherland) in a sealed compartment. The coil is located with its cylindrical axis vertically within a nylon panel enclosure on which petri dishes can be placed for microscope imaging. The coil and core were potted in high-temperature epoxy (Devcon, Milpitas, CA). The upper rim of coil windings and ferrite inside the compartment are 3 mm below the surface for placing petri dishes. The coil is electrically connected via two ¼-20” bolts to the cable, and both the coil and the ferrite compartment are cooled via chilled circulation attached via metal quick-disconnect fitting (CPC Colder, St. Paul, MN, USA). The coil’s electrical characteristics were measured in four-wire configuration with a bench LCR/ESR meter (Model 889A, B&K Precision Corporation, USA).

### 2.5. Control strategy

A SoC with field-programmable gate array (FPGA) and ARM Cortex A9 microcontroller (Xilinx Zynq 7000, sbRIO-9627, National Instrument, Austin, TX, USA) controls the transistor banks and the contactors. The SoC connects to a PC with a graphic user interface (LabVIEW, National Instruments, Austin, TX, USA) through an ethernet–USB adapter. The user interface selects as well as sets the stimulation channel and controls the output’s pulse duration, interpulse interval, stimulation frequency, and amplitude via pulse-width modulation (Figure 4). It can also program a sequence for repeated cycling of different channels in any specific order. The FPGA part of the SoC performs all transistor switching, for which we clocked it at 100 MHz to refine the time resolution so that the driver circuit can hit the resonant frequency close enough.

**Figure 4.**
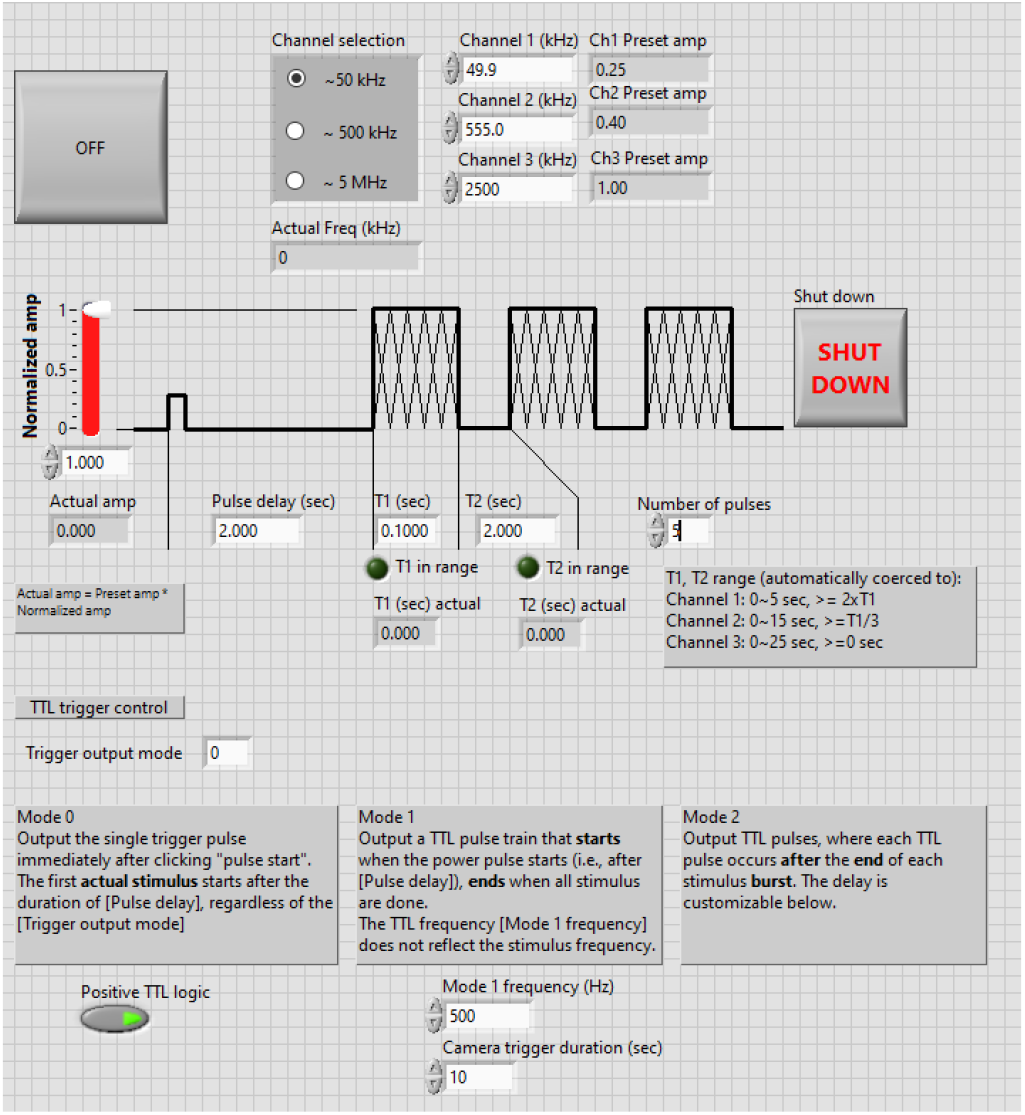
LabView graphic user interface for controlling the output.

### 2.6. System integration

All electronics components were placed in a standard 4 height-unit 19-inch vented rack (Hammond Manufacturing, Guelph, Canada), with the aluminum enclosure acting as shielding to limit RF noise leaking to the nearby equipment. Heat sinks were added on the electronics boards to increase thermal capacity and surface area and each circuit module had two fans, one above and one below, for cooling (Figure 5, left). A 12 V AC-DC adapter powers the FPGA and fans. Five 36 V 1 kW DC power supplies (Mean Well USA, Fremont, CA, USA) were used to power the electronics, two of which were placed on the bottom level of the rack. The three additional 1 kW power supplies were placed in a 2 height-unit 19-inch vented rack and this auxiliary power unit was connected to support the main system’s power supplies in parallel (Figure 5, right). A custom power cable was prepared to power the system via 3-phase 240 V NEMA15-30 outlets.

**Figure 5.**
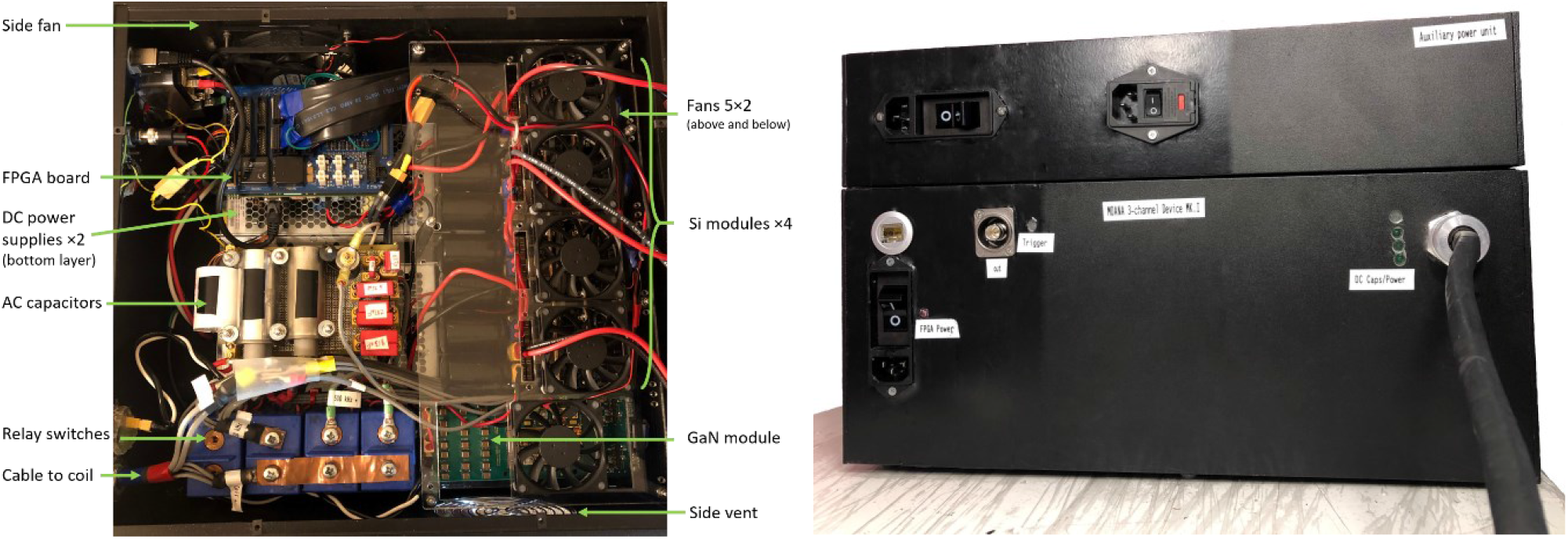
Left: the layout of the integrated system within the main unit. Right: The front panels of the main and auxiliary unit.

Waterjet cutting (Protomax, Omax Corp., Kent, WA, USA) was utilized to modify the front and side panels of the rack. Electric power was connected via power entry modules with circuit breakers (Schurter AG, Lucerne, Switzerland). The cable feeding the coil was connected to the relays via a cable gland. The FPGA board was connected externally to the PC via an ethernet port and provided a trigger output via a BNC connector to external equipment, such as a microscope camera for recording cellular response of the magnetogenetic stimulation and synchronizing the recording with the stimulation. The trigger (Figure 5, right) outputs a train of rectangular pulses that can be programmed with a specific frequency, pulse number, and delay time until the stimulation output, corresponding to the frames per seconds, total frames, and time for imaging baseline activity. Finally, LED status lights were installed on the front panel, and a fan was installed on the side panel.

### 2.7. Testing set-up and measurements

We performed all electrical measurements with a 500 MHz high-bandwidth oscilloscope (Tektronix MDO3054, Beaverton, OR, USA). High-voltage differential probes (Tektronix THDP0100) served to measure the coil voltage, whereas Rogowski current transducers (CWT 60B & CWT MiniHF 1B, Power Electronic Measurement Ltd., Nottingham UK, for Ch 1 & Ch 2 and Ch 2 & Ch 3, respectively) detected the coil current. To measure the magnetic field generated by the coil, we designed and manufactured a magnetic flux test coil of ten turns by winding 30-gauge magnetic wire around a 3D-printed polylactic acid cylindrical former of 1 cm diameter (Ultimaker B.V., Utrecht, Netherlands); the other end of the wire was twisted together and connected to the oscilloscope via a BNC connector. The probe measures the voltage induced by the changing magnetic flux of the alternating magnetic field so that the field strength can be easily derived in either the time or frequency domain.

Resonance frequencies for each channel were confirmed by varying the output frequency of the electronics circuit and maximizing the measured coil current/voltage or induced voltage of the magnetic field probe. Once the resonance frequency was recorded, it was not changed any more during operation.

We tested sequenced cycling through the three channels with the following settings: Ch 1, 49.9 kHz, 90 mT; Ch 2, 549 kHz, 13.7 mT; Ch 3, 2.5 MHz, 3.9 mT (see Section 3.2 for results on frequency and amplitude combinations of the channels). The channels were cycled in the order from lowest to highest frequency with equal pulse duration of 3 s and two inter-pulse intervals of 17 s and 7 s were tested, corresponding to a duty cycle and coil current of 15% with 88 A and 30% with 125 A, respectively. Temperatures of the coil, cable, and electrical and electronics components within the rack were measured using a handheld infrared thermometer (Fluke Corporation, Everett, WA, USA).

Data analysis was performed in MATLAB (R2019b, The MathWorks, Natick, MA). To calculate the magnetic field, the induced voltage was detrended to remove any DC offset, filtered above 50 MHz, and integrated.

### 2.8. Nanoparticle synthesis and specific absorption rate measurement

Three types of iron oxide nanoparticles with different coercivity were developed to form three magnetothermal channels. The 15 nm cobalt-doped iron oxide (Fe_2.35_Co_0.65_O_4_) nanoparticles, 19 nm iron oxide (Fe_3_O_4_) nanoparticle, and 10 nm manganese-doped iron oxide (Fe_2.43_Mn_0.57_O_4_) nanoparticles generate a large amount of heat respectively in a low-frequency high-amplitude magnetic field (Ch 1), a medium-frequency medium-amplitude field (Ch 2), and a high-frequency low-amplitude field (Ch 3). The synthesis, coating, and functionalization of these nanoparticles followed previous methods (Tong *et al.* 2017; Sebesta *et al.* 2021).

Briefly, magnetite nanocrystals of 4 nm diameter were first synthesized by thermal decomposition of iron acetylacetonate in a mixture of oleic acid and benzyl ether and grown to 15 nm diameter by controllable seed-mediated growth in a mixture of iron acetylacetonate (Fe(C_5_H_7_O_2_)_3_), oleic acid, and benzyl ether. Then, the nanocrystals were coated with a copolymers of phospholipids and polyethylene glycol (DSPE-PEG2K) using a dual-solvent exchange method (Tong *et al.* 2011). The cobalt-doped nanoparticles were synthesized from multiple seed-mediated growth reactions through thermal decomposition with 5 nm iron oxide cores in 2 mmol CoCl_2_, 4 mmol Fe(C_5_H_7_O_2_)_3_, 25 mmol oleic acid, using 60 mL benzyl ether as a solvent. 7 nm manganese-doped seeds were synthesized with a mixture of 5 mmol MnCl_2_, 10 mmol Fe(C_5_H_7_O_2_)_3_, 50 mmol 1,2-tetradecanediol, 60 mmol oleic acid, 60 mmol oleylamine and 60 ml benzyl ether. The seeds were then grown to 10 nm with a seed-mediated growth procedure. The reacting solution was heated to 120 °C for 30 minutes under a constant argon flow, then to 200 °C for 2 hours and finally to reflux at 300 °C for 30 minutes. The product was purified through several acetone washes. The nanoparticle sizes were determined by HC TEM. Nanoparticles were then coated with DSPE-PEG2K by mixing the nanoparticles with PEG and adding DMSO. The reaction was then evaporated and transferred to water by a drop-wise addition of water and removal of the remaining DMSO by was done by centrifugation and ultracentrifugation.

We performed transmission electron microscopy (TEM), superconducting quantum interference device (SQUID), and power X-ray diffraction (XRD) measurements to quantify the size distribution, magnetic properties, and the crystal structure of the magnetite nanocrystals, respectively. The hydrodynamic size of conjugated nanoparticles was subsequently examined by dynamic light scattering. The samples for TEM measurements, both HC TEM and TITAN TEM, were prepared by diluting the samples and placing them in carbon-film grids. XRD samples were prepared by drying the nanoparticles under an argon flow and then pulverizing the resulting powder. SQUID measurements were performed with coated samples by fixing the nanoparticles with calcium hemisulfate and enclosing them within a capsule to prevent movement. Doping percentages were determined by inductively coupled plasma mass spectrometry and comparing the samples to the corresponding standard curves of iron, cobalt, and manganese.

The nanoparticle samples were placed on the surface of the coil and the heating was measured using an infrared thermal probe (Lumasense Luxtron 812 and STF-2M Probe). The specific absorption rate (SAR) is calculated as

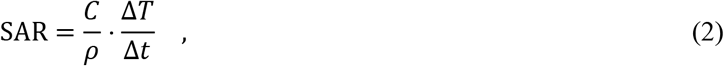

where *C* is the specific heat capacity of the media (4180 J·kg^−1^·K^−1^), Δ*T* is the temperature change during stimulation averaged over 3 stimulations, Δ*t* is the stimulation time, and *ρ* is sample density measured by total metal concentration. Iron oxide nanoclusters were recorded at 4.33 mg_metal/mL_, cobalt-doped nanoparticles were recorded at 9.58 mg_metal/mL_, and manganese-doped nanoparticles were recorded at 28.18 mg_metal/mL_.

## 3. Results

### 3.1. Electric characteristics of the coil

The high-flux coil had a measured inductance of *L*_coil_ = 3.14 μH and resistance of *R*_coil_ = 6.5 mΩ at 10 kHz and the cable added 0.13 μH and 2.5 mΩ. Using the custom-made magnetic field probe, the magnetic field distribution was mapped within the petri dish chamber. The coil demonstrates a relative uniform distribution over the central 1 cm region (Figure 6), with a gain of 0.134 mT/A and 0.108 mT/A at a height of 3 mm and 6 mm above the center cylinder, respectively, which corresponds to distances of 6 mm and 9 mm above the ferrite and top coil winding.

**Figure 6.**
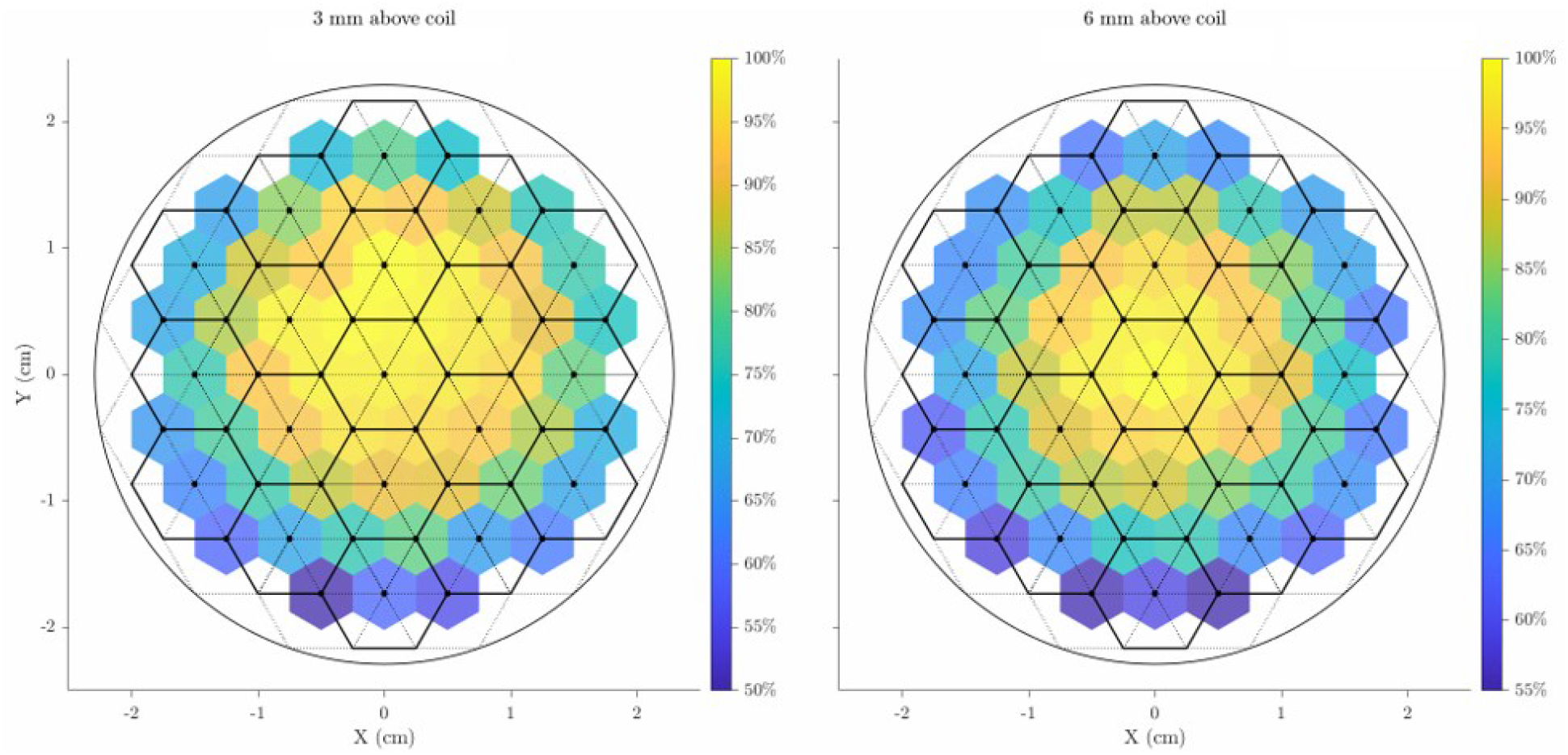
The normalized magnetic field distribution, 3 mm above the surface of the center cylinder (left) and 6 mm above the center cylinder.

### 3.2. Customization of resonance frequency and field measurement

While the software ensures hitting the resonant frequencies, our customizable capacitor arrays allow further adjustment on the resonant frequencies (Table 3 and Figure 8). Ch 1 resonated at 49.7 kHz and 61.5 kHz with 2 μF and 3 μF capacitors, respectively. Ch 2’s output has 46 frequencies in the 380–580 kHz frequency range as allowed by the FPGA clock, and the system was characterized with approximately 25 kHz steps between 400–575°kHz. Ch 3 had 15 frequencies within the 2–5 MHz range allowable by the FPGA’s clock. Among these options, we finalized our system at 49.7 kHz, 549 kHz, and 2.5 MHz, or the user can easily change the frequency for each channel using the capacitor combinations in Table 3.

**Table 3.** List of frequencies and capacitors for each channel

| Channel | Frequency (kHz) | Capacitors<sup>#</sup> (nF) | Channel | Frequency (MHz) | Capacitors<sup>#</sup> (pF) |
| --- | --- | --- | --- | --- | --- |
| Ch 1 | 49.7 | $1000 \times 2$ | Ch 3 | 2.50 | $470 + 150 \times 2$ |
| | 61.5 | $1000 \times 3$ | | 2.63 | $470 + 100 \times 2$ |
| Ch 2 | 400 | $22 + 10 + 6.8 \times 2$ | | 2.78 | $470 + 100$ |
| | 424 | $15 \times 2 + 10$ | | 2.94 | 470 |
| | 450 | $15 + 6.8 \times 3$ | | 3.13 | $150 + 100 \times 2$ |
| | 476 | $15 + 10 + 6.8$ | | 3.33 | $150 \times 2$ |
| | 500 | $10 \times 2 + 4.7 \times 2$ | | 3.57 | $100 \times 2$ |
| | 526 | $4.7 \times 5 + 3.3$ | | 3.85 | 100 |
| | 549 | $6.8 + 4.7 \times 3 + 3.3$ | | 4.17 | $100 \parallel 100$ |
| | 575 | $10 + 6.8 + 4.7$ | | 4.55 | 0 |

<sup>#</sup> Capacitors for Ch 1 have 10% tolerance, and those for Ch 2 and Ch 3 have 5% tolerance. The Ch 3 capacitors are in series with a 10–100 pF variable capacitor, which is tuned to reach resonance for each frequency tested.

The measured magnetic field amplitude (Figure 7) shows that the target field strength can be achieved for each channel. However, due to the higher impedance (Figure 8) and losses with increased frequency, the achievable magnetic field decrease for higher frequencies and the envisioned target of 1.25 mT was only achieved for frequencies between ≤4 MHz. The fitted coil impedance (Figure 8) of 3.12 μH and 83.3 mΩ matched the inductance measurement of the LCR meter, whereas the increased resistance of the coil can be attributed to the skin effect at higher frequency.

**Figure 7.**
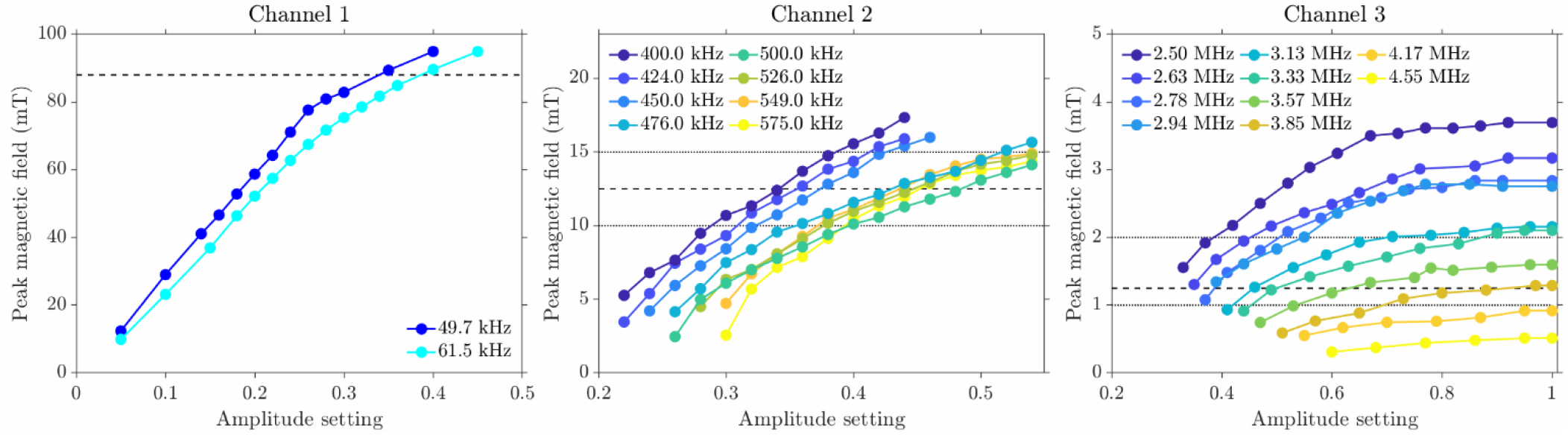
The output characteristics of the three channels. The amplitude setting determines the pulse-width modulation of the control voltage on the gates of the transistors. The dashed line shows the nominal target field strength for each channel, with the dotted lines indicating the upper (for Ch 2 and Ch 3 channel only) and lower ranges of the target.

**Figure 8.**
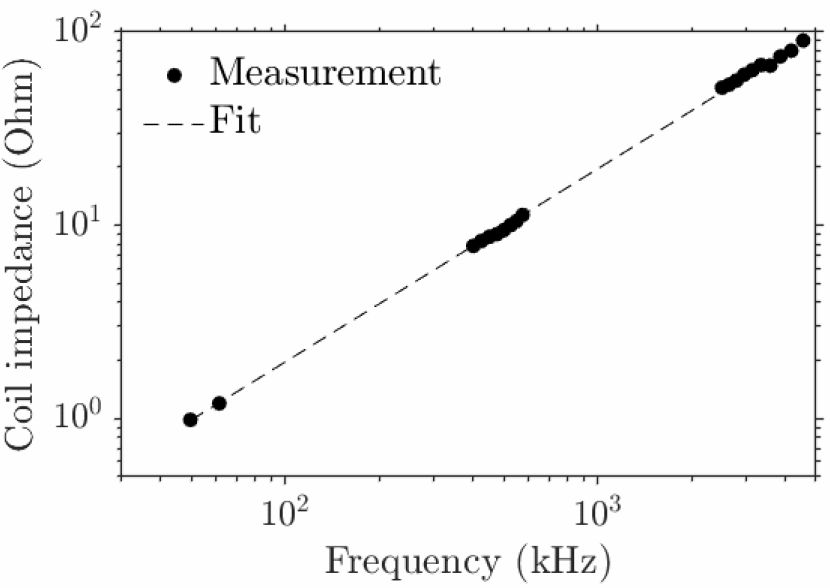
The coil impedance as a function of frequency.

### 3.3. Channel switching time, time for oscillation to reach steady state and decay to base line

After the transistors start switching, the L–C circuit begins oscillation with increasing amplitude until reaching steady state. As the power electronics allow pulse-width modulation, the envelope of this transient can be fitted with an exponential rise and decay at the beginning and end of the stimulation pulse, with time constant inversely proportional to the frequency. Ch 1’s time constant was approximately 0.2 ms, requiring about 1 ms to reach steady state or for the oscillation to stop (Figure 9, top); for Ch 2 and Ch 3, the transient’s durations were 10- and 100-times shorter, respectively, and therefore negligible compared to the minimum pulse duration anticipated (10 ms).

**Figure 9.**
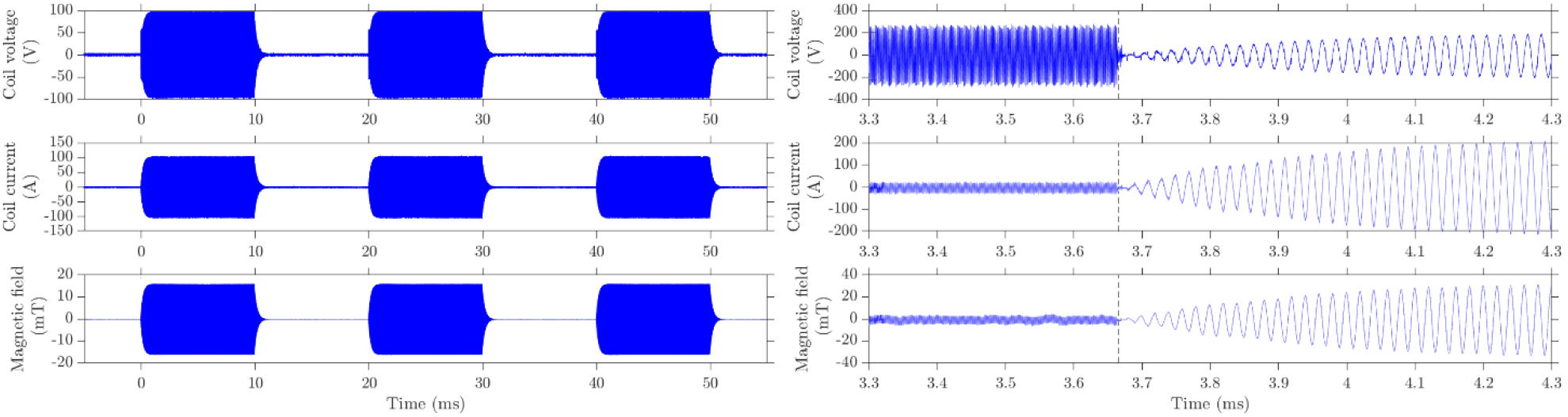
The coil voltage, current, and magnetic field 3 mm above the center of the coil. Left: Ch 1 switching on and off with 10 ms pulse duration and 10 ms interpulse intervals. Right: Switching from Ch 2 to Ch 3, dashed vertical line indicating switch time.

Within each channel, the pulse can be turned on and off with sub-millisecond speed. Between channels, the switching was slower for the contactors to close or break contact. Also, for safety reasons, a short delay was inserted in the control program when switching between different channels to avoid the high current from one channel and voltage from one capacitor bank being directly applied to another one. The switching between channels therefore can be achieved no slower than 1 ms (Figure 9, bottom).

### 3.4. Operation temperature and stability

Ch 1 and Ch 2 are thermally limited by temperature rise in the Si transistors for the presented current levels, the series AC capacitors, the cable and the coil, and may run single pulses up to 10 s and 30 s duration, respectively. Ch 3 can achieve continuous operation. To test the stability of the system, a pulse train cycling through the three channels was output with equal pulse duration and interpulse intervals at two different duty cycles. The system was able to continuously operate for two hours, with the temperature reaching a steady cycle around 20 minutes. Besides the cable, which reached 90 °C for the 30% duty cycle case, most components had moderate operating temperature (Figure 10) and were well within their specifications. According to these experimental results, it is advised to budget more on the cable since it bottlenecks our thermal performance, and thus likely to consume a large share of the supplied power.

**Figure 10.**
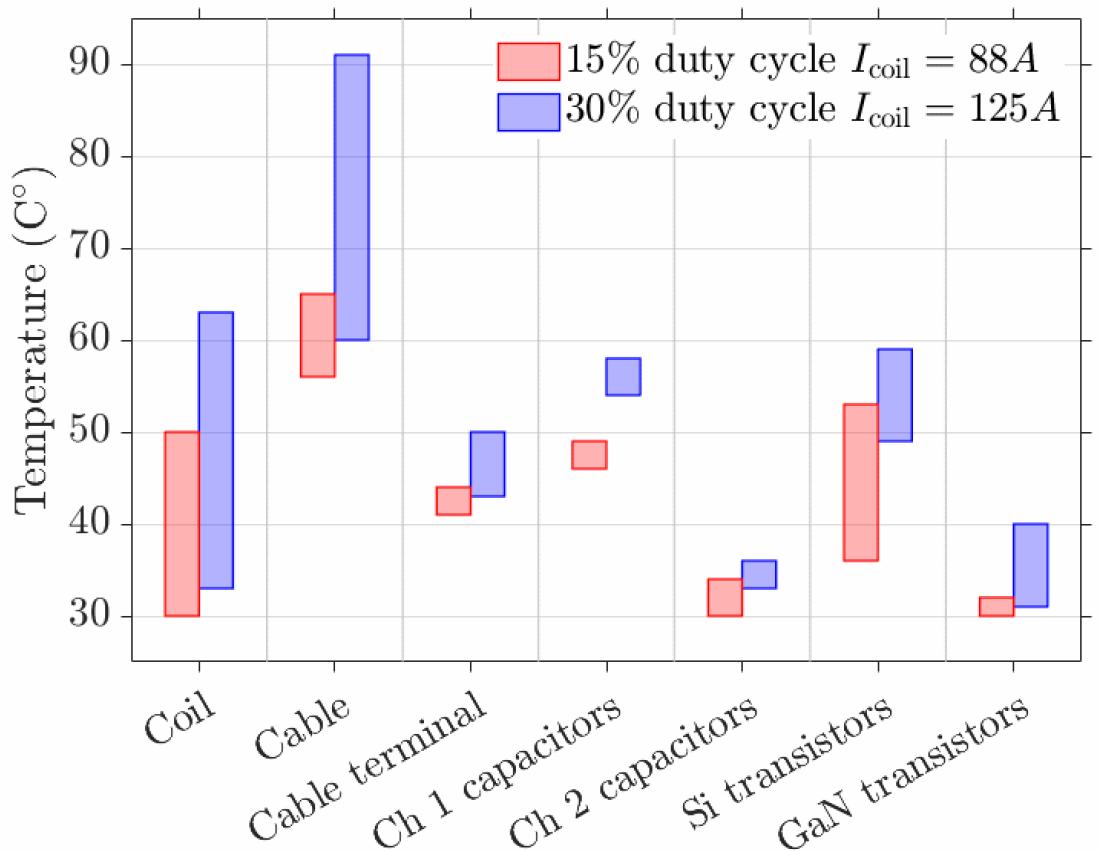
Temperature of different components of the system during two hours of continuous operation. The range indicates the maximum and minimum temperature during the cycle of three low, medium, and high frequency pulses. The maximum was recorded at the end of the Ch 1 pulse, except for the Ch 2 capacitors and GaN transistors, for which the maximum was recorded at the end of the Ch 2 and Ch 3 pulses, respectively.

### 3.5. Specific absorption rate of nanoparticles and channel selectivity

The heating of the three types of nanoparticles over 3 s or 15 s magnetic field stimulation in all three channels is shown in Figure 11, with the respective SAR summarized in Table 4. When stimulation time is limited to 3 s for Ch 1 and 15 s for Ch 2 and 3, the maximum temperature change showed a selectivity of ∼9× for 15 nm Cobalt doped NPs in Ch 1, ∼2.3× for 19 nm iron-oxide NPs in Ch 2, and ∼3.7× for 10 nm Mn-doped NPs in Ch 3, thus indicating three distinct channels available for magnetothermal heating. By adjusting the concentration of the nanoparticles and the corresponding stimulation time duration, three samples within the same magnetic field could be separately activated (Figure 12).

**Table 4.** Specific absorption rate of the 3 types of nanoparticles measured in the three channels.

| SAR ($\text{W}\cdot\text{g}^{-1}$) | Channel 1 | Channel 2 | Channel 3 |
| --- | --- | --- | --- |
| 15 nm Co-doped iron oxide | 847.57 | 18.44 | 4.80 |
| 19 nm iron oxide | 151.63 | 123.48 | 54.43 |
| 10 nm Mn-doped iron oxide | 12.31 | 3.47 | 12.96 |

**Figure 11.**
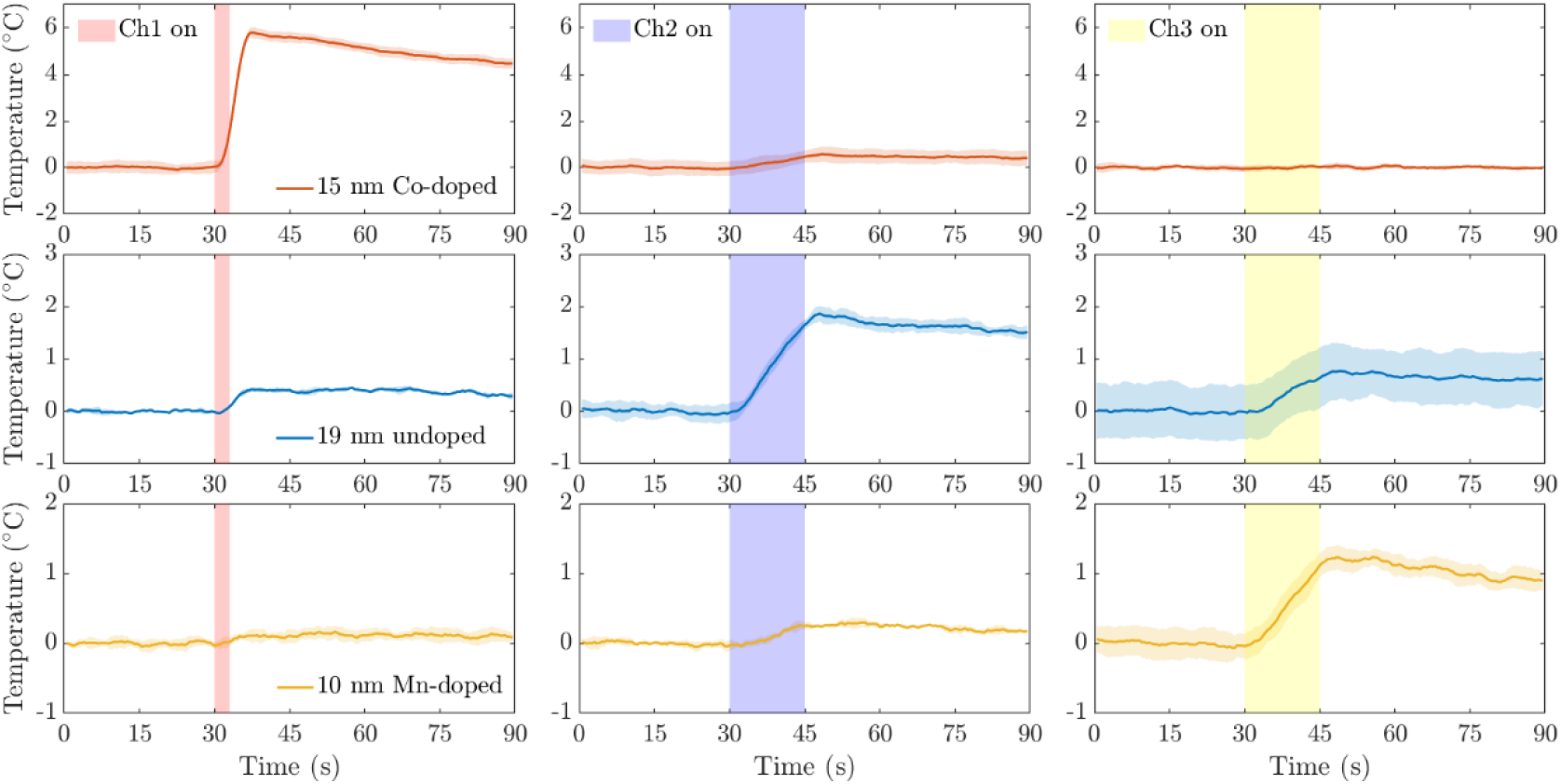
Thermal response of 15 nm cobalt-doped iron oxide nanoparticles (red traces, top row), 19 nm iron oxide nanoparticles (blue, middle row), and 10 nm manganese-doped iron oxide nanoparticles (yellow, bottom row) when exposed to a magnetic field of Ch 1 (left column, 49.9 kHz; 80 mT), Ch 2 (center column, 555 kHz; 12 mT), or Ch 3 (right column, 2.227 MHz, 3.8 mT).

**Figure 12.**
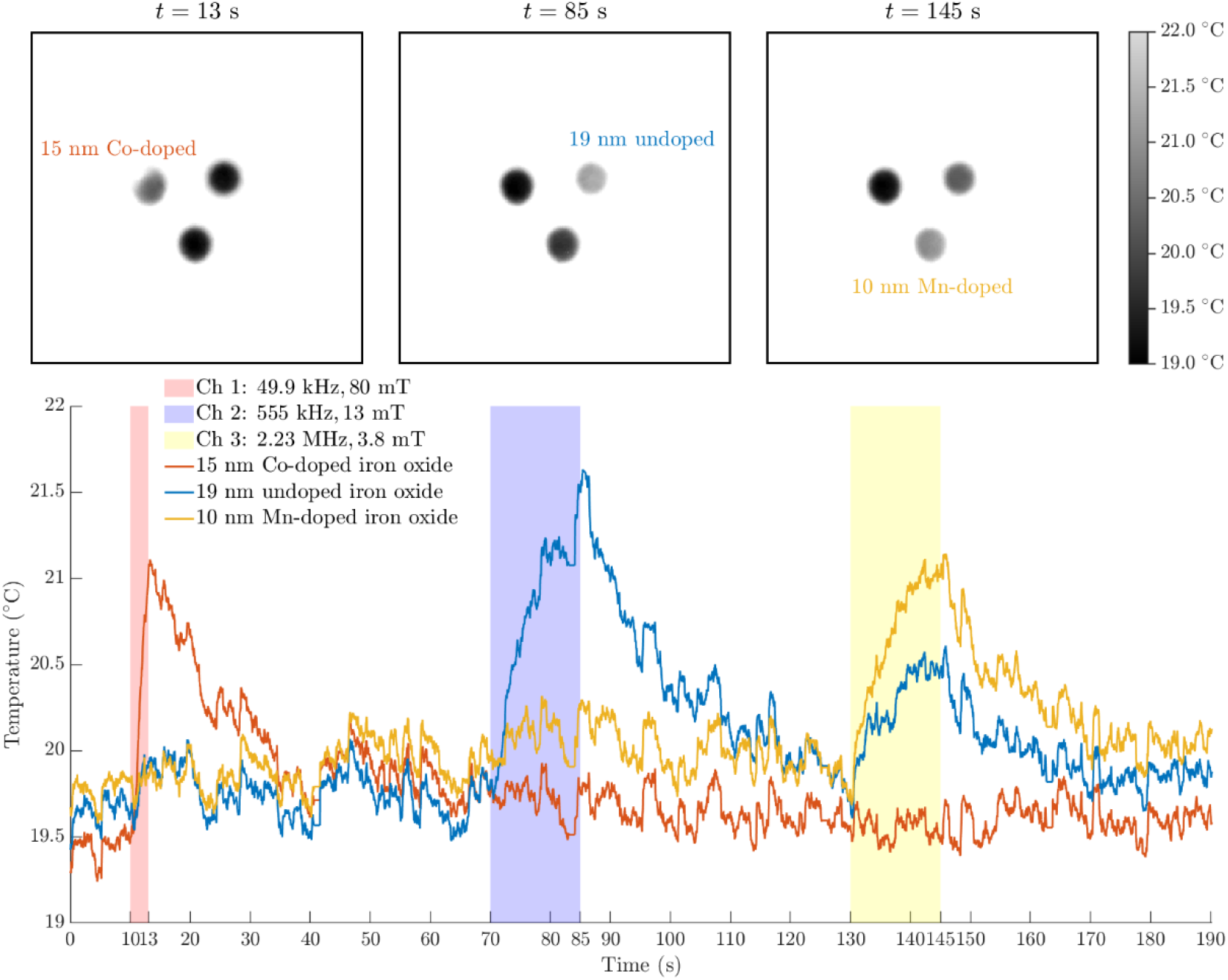
Thermal imaging recorded with infrared camera (FLIR A700) of 10 μL drops of each sample at concentrations adjusted to generate similar amounts of total heat. Channel 1 shows selective heating of 15 nm Co-doped iron oxide (∼8 mg/mL), whereas Ch 2 shows selective heating of 19 nm iron oxide (∼3 mg/mL) and channel 3 shows selective heating of 10 nm Mn-doped iron oxide (∼22 mg/mL).

## 4. Discussion

### 4.1. Power electronics design

In this study, we designed and implemented a power electronics system to drive via series resonance a commercially available coil at different frequencies spanning from low kilohertz to megahertz, thus forming distinct channels for multiplexed magnetogenetic stimulation, with fast and flexible switching between the channels. The key innovation is to adopt a hybrid approach using two types of transistors, optimized for the different frequency and current levels required, and high voltage contactors for switching channels with millisecond speed. The Si transistors provide high current at the lower-frequency ranges, whereas the GaN transistors covered the highest frequencies, where the current magnitude is lower.

The system can select from various capacitor banks and temporarily connect them to the coil via contactors. In contrast to the transistors in the power electronic circuit, the contactors have to both conduct high currents when closed—at least for the low-frequency range—and block high voltage when opened. The switching speed of channels is primarily determined by the mechanical speed of the contactors. Such a combination is currently challenging in electronics. The achieved millisecond channel switching is sufficient for current magnetogenetic applications that typically have neural response time on the order of at least hundreds of milliseconds and up to several seconds (Huang *et al.* 2010; Chen *et al.* 2015; Munshi *et al.* 2017). Unlike several previous systems in which the capacitor bank is located next to the coil and sometimes an integrated component, our system integrates the capacitor banks with the power electronics unit and connects them to the coil via a relatively flexible and long cable. This separation of source and magnetic coil enables the placement of the coil under a microscope. Furthermore, it allows better combination with other units of typical magnetogenetic systems, such as temperature and humidity control as well as perfusion while having the power system placed far from the imaging set-up, reducing electronic noise and vibration. The use of Litz wire reduces high-frequency losses and the cable’s terminal design is capable of interfacing with other available coils, for example the commercial nanoTherics magneTherm™ coil (Drayton *et al.* 2017), which we tested during the prototyping phase.

### 4.2. Nanoparticle heating

Heating can be improved and potentially further multiplexed with the inclusion of multicore nanoclusters which enable higher heating rates for particles with lower anisotropy (Xiao *et al.* 2020; Sebesta *et al.* 2021). However, this study demonstrates a proof of concept for 3 channels by modifying the magnetic anisotropy of single domain nanoparticles via doping and selectively adjusting the frequency and field strength of AMF stimulations to pair specifically with the anisotropy of those particles. As the anisotropy is lowered by the inclusion of manganese in the crystal structure, Mn-doped nanoparticles demonstrate limited heating rates through hysteretic heating which may have limited use for biological applications but show promise for a 3^rd^ multiplexed channel. Further optimization of doping concentrations and tuning of AMF may improve selectivity and performance for each channel which can be better targeted with the use of custom AC magnetometers (Moon *et al.* 2020).

### 4.3. Limitations

The system’s long-term operation was mostly restricted due to the thermal limitations of several components. For single pulses under low repetition rate, the limiting factors determining the maximum pulse duration for the Ch 1 and Ch 2 channels were the Si power electronics and the AC capacitors, whereas the coil does not experience significant heating. On the other hand, the MHz channel can run continuously. In addition to the Ch 1 and Ch 2 electronic components, repetitive pulsing needs to consider the cable and coil to avoid thermal stimulation of the neuronal targets.

The contactors allowed fast switching between the channels with different order-of-magnitude currents. Furthermore, the clicking of the contactors closing and breaking contact is clearly audible, despite them being located on an elevated platform inside the rack unit. Although this noise was tolerable and nowhere near hazardous levels to the experimental operator, its presence and associated auditory activation could be of concern when stimulating animals with intact auditory system, which might lead to well-synchronized indirect brain stimulation as known from other paradigms (Zaehle *et al.* 2007; Goetz *et al.* 2015).

### 4.4. Future direction and next generation system

The system scales well and allows modular expansion of capacitor banks and the transistor circuits to increase the current capabilities as well as the number of channels. The GaN transistors readily allow driving frequencies above 50 MHz. Thus, this system in combination with a coil design optimized for higher frequency intends to help find magnetic nanoparticles and proteins for higher frequency ranges to further increase multiplexing capabilities.

## 5. Conclusions

We developed a multi-channel power electronics system and magnetic nanoparticles for frequency-multiplexed magnetogenetic neurostimulation with three channels. The system spanned a wide frequency range and showed millisecond switching speed between channels as well as thermal stability for hour-long operation. We demonstrated selective heating of magnetic particles in all three channels by using a combination of the duration of each channel’s stimulation time and the concentration of the nanoparticles. This system will enable experimental studies of magnetogenetic neurostimulation with improved selectively, speed, and flexibility.

## Acknowledgment

This research was developed with funding from the Defense Advanced Research Projects Agency (DARPA) of the United States of America, Contract No. N66001-19-C-4020. The views, opinions and/or findings expressed are those of the authors and should not be interpreted as representing the official views or policies of the Department of Defense or the U.S. Government. The authors thank Dr. Zhiyong Zeng, Dr. Lari M. Koponen, and Rena Hamdan for helpful discussion and technical support. Waterjet cutting and 3D printing was performed in the Innovation Co-Lab of Duke University. Preliminary results were presented at Neural Interfaces 2021: The NANS-NIC Joint Meeting (June 2021, online).

